# Speech categorization is better described by induced rather than evoked neural activity

**DOI:** 10.1101/2020.10.20.347526

**Authors:** Md Sultan Mahmud, Mohammed Yeasin, Gavin M. Bidelman

**Affiliations:** Department of Electrical and Computer Engineering, University of Memphis, Memphis, TN 38152, USA; Institute for Intelligent Systems, University of Memphis, Memphis, TN 38152, USA; School of Communication Sciences & Disorders, University of Memphis, Memphis, TN 38152, USA; University of Tennessee Health Sciences Center, Department of Anatomy and Neurobiology, Memphis, TN 38103, USA

**Keywords:** Categorical perception, time-frequency analysis, induced oscillations, gamma-band activity, machine learning, support vector machine (SVM)

## Abstract

Categorical perception (CP) describes how the human brain categorizes speech despite inherent acoustic variability. We examined neural correlates of CP in both evoked and induced EEG activity to evaluate which mode best describes the process of speech categorization. Using source reconstructed EEG, we used band-specific evoked and induced neural activity to build parameter optimized support vector machine (SVMs) model to assess how well listeners’ speech categorization could be decoded via whole-brain and hemisphere-specific responses. We found whole-brain evoked β-band activity decoded prototypical from ambiguous speech sounds with ~70% accuracy. However, induced γ-band oscillations showed better decoding of speech categories with ~95% accuracy compared to evoked β-band activity (~70% accuracy). Induced high frequency (γ-band) oscillations dominated CP decoding in the left hemisphere, whereas lower frequency (θ-band) dominated decoding in the right hemisphere. Moreover, feature selection identified 14 brain regions carrying induced activity and 22 regions of evoked activity that were most salient in describing category-level speech representations. Among the areas and neural regimes explored, we found that induced γ-band modulations were most strongly associated with listeners’ behavioral CP. Our data suggest that the category-level organization of speech is dominated by relatively high frequency induced brain rhythms.

## I. INTRODUCTION

The human brain classifies diverse acoustic information into smaller, more meaningful groupings (Bidelman and Walker, 2017), a process known as categorical perception (CP). CP plays a critical role in auditory perception, speech acquisition, and language processing. Brain imaging studies have shown that neural responses elicited by prototypical speech sounds (i.e., those heard with a strong phonetic category) differentially engage Heschl’s (HG) and inferior frontal (IFG) gyrus compared to ambiguous speech (Bidelman et al., 2013; Bidelman and Lee, 2015; Bidelman and Walker, 2017). Previous studies also demonstrate that the N1 and P2 waves of the event-related potentials (ERPs) are highly sensitive to speech perception and correlate with CP (Alain, 2007; Bidelman et al., 2013; Mankel et al., 2020). These studies demonstrate that evoked activity in the time domain provides a neural correlate of speech categorization. However, ERP studies do not reveal how induced brain activity (so-called neural oscillations) might contribute to this process.

The electroencephalogram (EEG) can be divided into evoked (i.e., phase-locked) and induced (i.e., non-phase locked) responses that vary in a frequency-specific manner (Shahin et al., 2009). Evoked responses are largely related to the stimulus, whereas induced responses are additionally linked to different perceptual and cognitive processes that emerge during task engagement. These later brain oscillations (neural rhythms) play an important role in perceptual and cognitive processes and reflect different aspects of speech perception. For example, low frequency [e.g., θ (4-8 Hz)] bands are associated with syllable segmentation (Luo and Poeppel, 2012) whereas α (9-13 Hz) band has been linked with attention (Klimesch, 2012) and speech intelligibility (Dimitrijevic et al., 2017). Several studies report listeners’ speech categorization efficiency varies in accordance with their underlying induced and evoked neural activity (Bidelman et al., 2013; Bidelman and Alain, 2015; Bidelman and Lee, 2015). For instance, Bidelman assessed correlations between ongoing neural activity (e.g., induced activity) and the slopes of listeners’ identification functions, reflecting the strength of their CP (Bidelman, 2017). Listeners were slower and varied in their classification of more category-ambiguous speech sounds, which covaried with increases in induced γ activity (Bidelman, 2017). Changes in β (14-30 Hz) power are also strongly associated with listeners’ speech identification skills (Bidelman, 2015). The β frequency band is linked with auditory template matching (Shahin et al., 2009) between stimuli and internalized representations kept in memory (Bashivan et al., 2014), whereas the higher γ frequency range (>30 Hz) is associated with auditory object construction (Tallon-Baudry and Bertrand, 1999) and local network synchronization (Giraud and Poeppel, 2012; Haenschel et al., 2000; Si et al., 2017).

Studies also demonstrate hemispheric asymmetries in neural oscillations. During syllable processing, there is a dominance of γ frequency activity in LH and θ frequency activity in RH (Giraud et al., 2007; Morillon et al., 2012). Other studies show that during speech perception and production, lower frequency bands (3-6 Hz) better correlate with behavioral reaction times than higher frequencies (20-50 Hz) (Yellamsetty and Bidelman, 2018). Moreover, induced γ-band correlates with speech discrimination and perceptual computations during acoustic encoding (Ou and Law, 2018), further suggesting it reflects a neural representation of speech above and beyond evoked activity alone.

Still, given the high dimensionality of EEG data, it remains unclear which frequency bands, brain regions, and “modes” of neural function (i.e., evoked vs. induced signaling) are most conducive to describing the neurobiology of speech categorization. To this end, the recent application of machine learning (ML) to neuroscience data might prove useful in identifying the most salient features of brain activity that predict human behaviors. ML is an entirely data-driven approach that “decodes” neural data with minimal assumptions on the nature of exact representation or where those representations emerge. Germane to the current study, ML has been successfully applied to decode the speed of listeners’ speech identification (Al-Fahad et al., 2020) and related receptive language brain networks (Mahmud et al., 2020) from multichannel EEGs.

Departing from previous hypothesis-driven studies (Bidelman, 2017; Bidelman and Alain, 2015; Bidelman and Walker, 2017), the present work used a comprehensive data-driven approach (i.e., stability selection and SVM classifiers) to investigate the neural mechanisms of speech categorization using whole-brain electrophysiological data. Our goals were to evaluate which neural regime [i.e., evoked (phase-synchronized ERP) vs. induced oscillations], frequency bands, and brain regions are most associated with CP using whole-brain activity via a data-driven approach. Based on prior work, we hypothesized that evoked and induced brain responses would both differentiate the degree to which speech sounds carry category-level information (i.e., prototypical vs. ambiguous sounds from an acoustic-phonetic continuum). However, we predicted induced activity would best distinguish category-level speech representations, suggesting a dominance of endogenous brain rhythms in describing the neural underpinnings of CP.

## II. MATERIALS & METHODS

### A. Participants

Forty-eight young (male: 15, female: 33; aged 18 to 33 years) participated in the study (Bidelman et al., 2020; Bidelman and Walker, 2017; Mankel et al., 2020). All participants had normal hearing sensitivity (i.e., <25 dB HL between 500-2000 Hz) and no previous history of neurological disease. Listeners were-right handed and had achieved a collegiate level of education. All participants were paid for their time and gave informed written consent in accordance with the declaration of Helsinki and a protocol approved by the Institutional Review Board at the University of Memphis.

### B. Stimuli & task

We used a synthetic five-step vowel token continuum to examine the most discriminating brain activity (i.e., evoked or induced activity) while categorizing prototypical vowel speech sounds from ambiguous speech (Bidelman et al., 2013). Speech spectrograms are represented in FIG. 1A. Each speech token was 100 ms, including 10 ms rise/fall to minimize the spectral splatter in the stimuli. Each speech token contained an identical voice fundamental frequency (F0), second (F2), and third formant (F3) frequencies (F0:150 Hz, F2: 1090 Hz, and F3:2350 Hz). The first formant (F1) was varied over five equidistance steps (430-730 Hz) to produce perceptual continuum from /u/ to /a/.

**FIG. 1.**
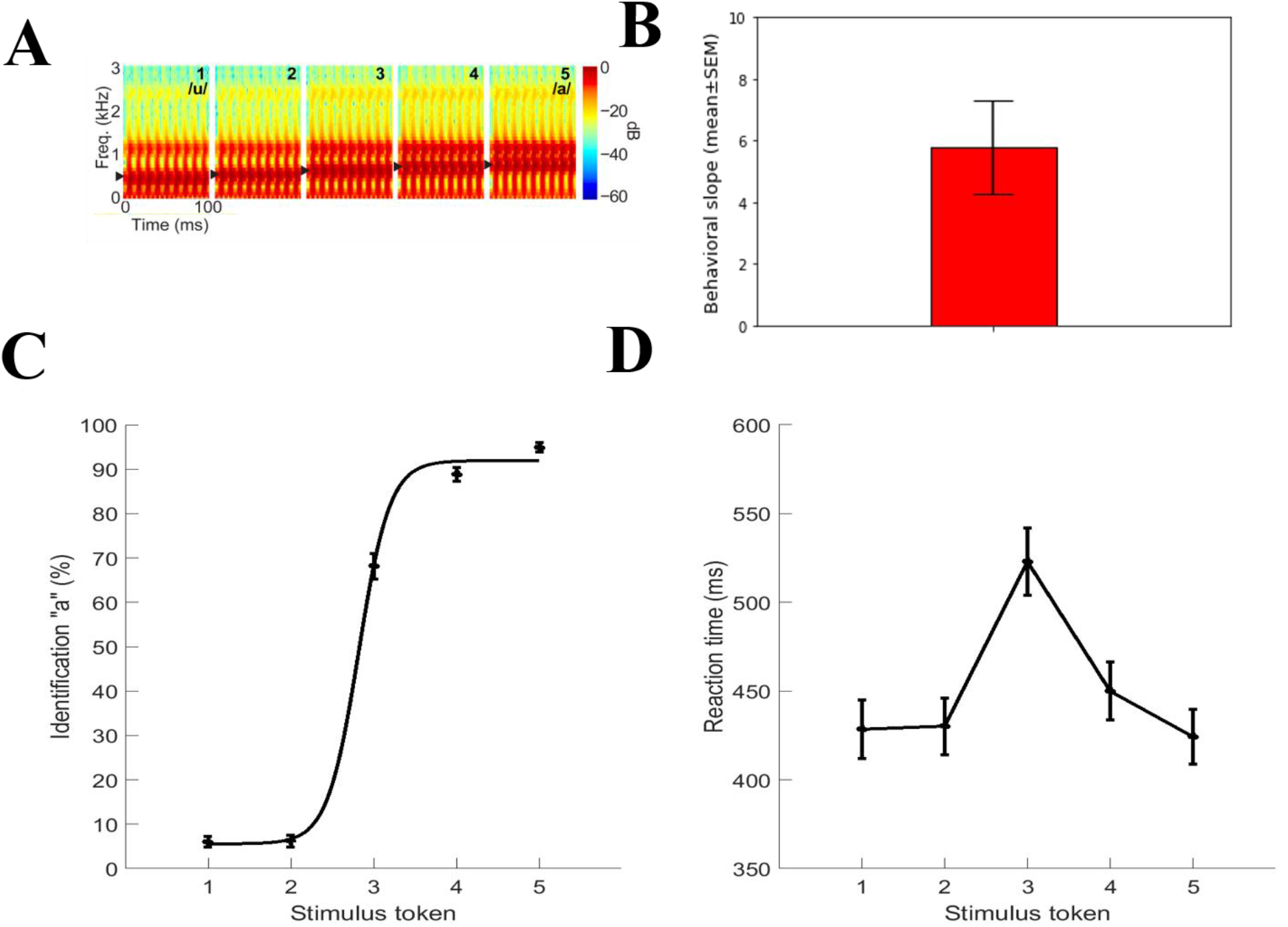
(Color online) Speech stimuli and behavioral results. A) Acoustic spectrograms of the speech continuum from */u/* and */a/*. B) Behavioral slope. C) Psychometric functions showing % “a” identification of each token. Listeners’ perception abruptly shifts near the continuum midpoint, reflecting a flip in perceived phonetic category (i.e., “u” to “a”). D) Reaction time (RT) for identifying each token. RTs are faster for prototype tokens (i.e., Tk1/5) and slow when categorizing ambiguous tokens at the continuum’s midpoint (i.e., Tk3). Errorbars = ±1 s.e.m.

Stimuli were delivered binaurally at an intensity of 83 dB SPL through earphones (ER 2; Etymotic Research). Participants heard each token 150-200 times presented in random order. They were asked to label each sound in a binary identification task (“/u/” or “/a/”) as fast and accurately as possible. Their response and reaction time were logged. The interstimulus interval (ISI) was jittered randomly between 400 and 600 ms with 20 ms step.

### C. EEG recordings and data pre-procedures

EEGs were recorded from 64 channels at standard 10-10 electrode locations on the scalp and digitized at 500 Hz using Neuroscan amplifiers (SynAmps RT). Subsequent preprocessing was conducted in the Curry 7 neuroimaging software suite, and customized routines coded in MATLAB. Ocular artifacts (e.g., eye-blinks) were corrected in the continuous EEG using principal component analysis (PCA) and then filtered (1-100 Hz; notched filtered 60 Hz). Cleaned EEGs were then epoched into single trials (−200 to 800 ms, where *t* = 0 was stimulus onset) and common average referenced. For details see in (Bidelman et al., 2020; Bidelman and Walker, 2017).

### D. EEG source localization

To disentangle the functional generators of CP-related EEG activity, we reconstructed the sources of the scalp recorded EEG by performing a distributed source analysis on single-trial data in Brainstorm software (Tadel et al., 2011). We used a realistic boundary element head model (BEM) volume conductor and standard low-resolution brain electromagnetic tomography (sLORETA) as the inverse solution within Brainstorm (Tadel et al., 2011). From each single-trial sLORETA volume, we extracted the time-courses within the 68 functional regions of interest (ROIs) across the left and right hemispheres defined by the Desikan-Killiany (DK) atlas (Desikan et al., 2006) (LH: 34 ROIs and RH: 34 ROIs). Single-trial data were baseline corrected to the epoch’s pre-stimulus interval (−200-0 ms).

Since we were interested in decoding prototypical (Tk1/5) from ambiguous speech (Tk3), we merged Tk1 and Tk5 responses since they reflect prototypical vowel categories (“u” vs. “a”). In contrast, Tk3 reflects a bistable percept—an ambiguous category listeners sometimes label as “u” or “a” (Bidelman et al., 2020; Bidelman and Walker, 2017; Mankel et al., 2020). To ensure an equal number of trials for prototypical and ambiguous stimuli, we considered 50% of the data from the merged samples.

### E. Time-frequency analysis

Time-frequency analysis was conducted via wavelet transform (Herrmann et al., 2014). First, we computed the event-related potential (ERP) using bootstrapping by randomly averaged over 100 trials with the replacement for each stimulus condition (e.g., Tk1/5 and Tk3) per subject and source ROI (e.g., 68 ROIs). We then applied the Morlet wavelet transform to the averaged data (i.e., ERP) with an increment step frequency 1 Hz from low to high frequency (e.g., 1 to 100 Hz), which provided only evoked frequency-specific activity (i.e., time- and phase-locked to stimulus onset). For computing induced activity, we performed a similar Morlet wavelet transform on a single-trial basis for each ROI, and then computed the absolute value of each trial spectrogram. We then averaged the resulting time-frequency decompositions (Herrmann et al., 2014), resulting in a spectral representation that contains total activity. To isolate induced responses, we subtracted the evoked activity from the total activity (Herrmann et al., 2014). We then extracted the different frequency band signals from evoked and induced activity time-frequency maps for each brain region (e.g., 68 ROIs). Example evoked and induced time-frequency maps from the primary auditory cortex [i.e., transverse temporal (TRANs)] are represented in FIG. 2.

**FIG. 2.**
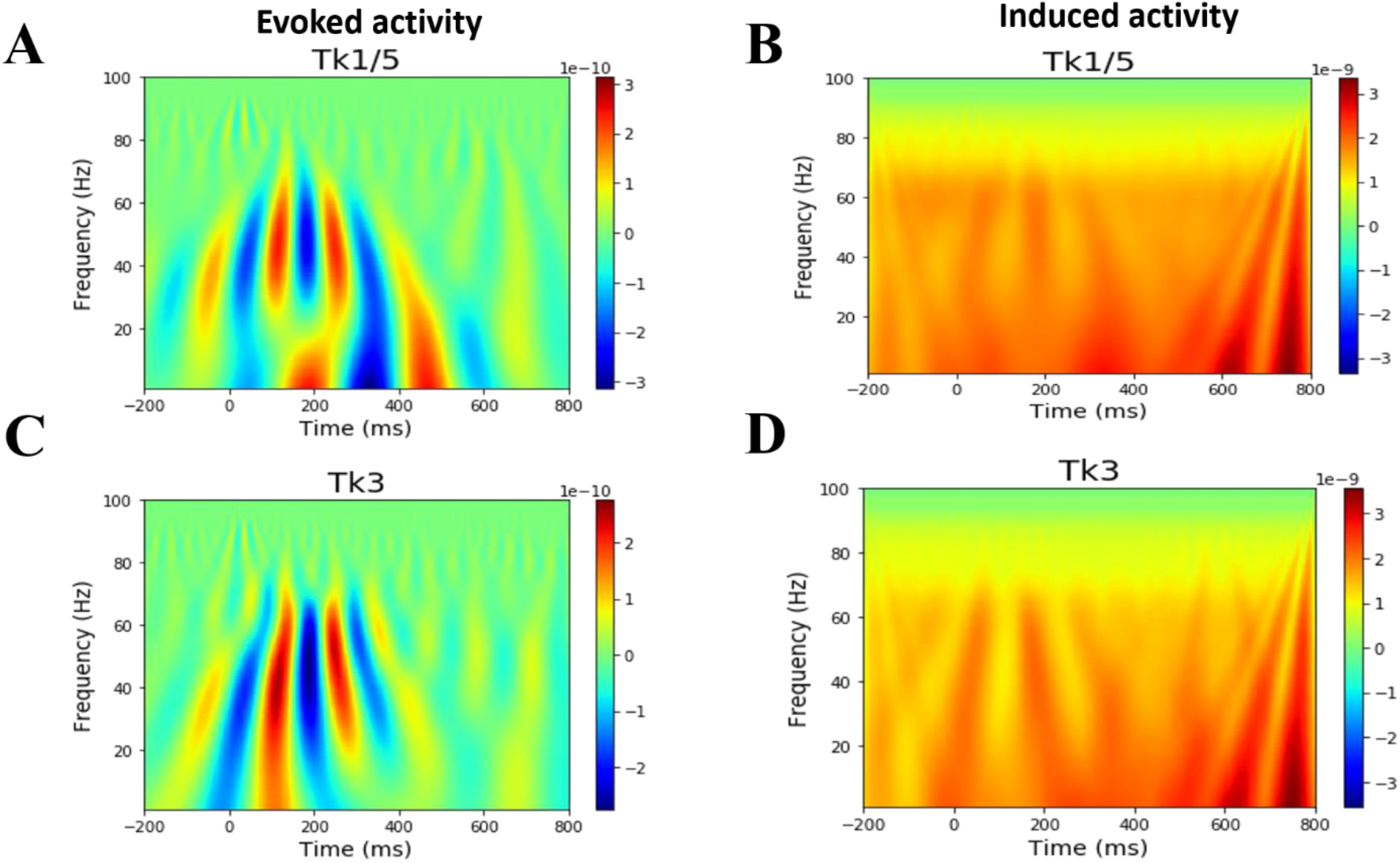
(Color online) Grand average neural oscillatory responses to prototypical vowel (e.g., Tk1/5 and ambiguous speech token (Tk3) A,C) Evoked activity for prototypical vs. ambiguous tokens. B, D) Induced activity for prototypical vs. ambiguous tokens. Primary auditory cortex (PAC) [lTRANS, left transverse temporal gyrus].

Spectral features of different bands (θ, α, β, and γ) were quantified as the mean power over the full epoch. We concatenated four frequency bands that resulted in 4*68=272 features for each response type (e.g., evoked vs. induced) per speech condition (Tk1/5 vs. Tk3). We separately (i.e., evoked and induced) submitted these individual frequency bands to the support vector machine (SVM) classifier and all concatenated features to stability selection to investigate which frequency bands and brain regions decode prototypical (e.g., Tk1/5) from ambiguous (Tk3) vowels. Features were z-scored prior to SVM to normalize them to a common range.

### F. SVM classification

We used the parameter optimized SVM that yields better classification performance with small sample sizes data (Mahmud et al., 2020). The tunable parameters (e.g., kernel, C, Ɣ) in the SVM model greatly affect the classification performance. We randomly split the data into training and test sets 80%, 20 %, respectively. During the training phase (i.e., using 80% data), we conducted a grid search approach with five-fold cross-validation, kernels = ‘RBF’, fine-tuned the *C*, and *γ* parameters to find the optimal values; so that the classifier can accurately distinguish prototypical vs. ambiguous speech (Tk1/5 vs.Tk3) in the test data that models have never seen. Once the models were trained, we selected the best model with the optimal value of *C*, and *γ* and predict the unseen test data (by providing the attributes but no class labels). Classification performance metrics (accuracy, F1-score, precision, and recall) were calculated using standard formulas. The optimal values of C and ɣ for different analysis scenarios are given in the Appendix.

### G. Stability selection to identify critical brain regions of CP

Our data comprised a large number (272 features) of spectral measurements for each stimulus condition of interest (e.g., Tk1/5 vs. Tk3) per brain regime (e.g., evoked and induced). We aimed to select a limited set of the most salient discriminating features via stability selection (Meinshausen and Bühlmann, 2010) (see details in the Appendix).

During the stability selection implementation, we considered a sample fraction = 0.75, number of resamples = 1000, and tolerance = 0.01 (Meinshausen and Bühlmann, 2010). In the Lasso algorithm, the feature scores were scaled between 0 to 1, where 0 is the lowest score (i.e., irrelevant feature) and 1 is the highest score (i.e., most salient or stable feature). We estimated the regularization parameter from the data using the least angle regression (LARs) algorithm. Over 1000 iterations, Randomized Lasso provided the overall feature scores (0~1) based on the number of times a variable was selected. We ranked stability scores to identify the most important, consistent, stable, and invariant features that could decode speech categories via the EEG. We submitted these ranked features and corresponding class labels to an SVM classifier with different stability thresholds and observed the model performance.

## III. RESULTS

### A. Behavioral results

Listeners’ behavioral identification (%) functions and reaction time (ms) for speech tokens categorization are illustrated in FIG. 1C and FIG. 1D, respectively. Responses abruptly shifted in speech identity (/u/ vs. /a/) near the midpoint of the continuum, reflecting a change in the perceived category. The behavioral speed of speech labeling (e.g., reaction time (RT)) was computed from listeners’ median response latency for a given condition across all trials. RTs outside of 250-2500 ms were deemed outliers and excluded from further analysis (Bidelman et al., 2013; Bidelman and Walker, 2017). For each continuum, individual identification scores were fit with a two-parameter sigmoid function; 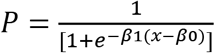, where *P* is the proportion of the trial identification as a function of a given vowel, *x* is the step number along the stimulus continuum, and *β0* and *β1* the location and slope of the logistic fit estimated using the nonlinear least-squares regression (Bidelman and Walker, 2017). The slope of listeners’ sigmoidal psychometric function reflects the strength of their CP (FIG. 1B).

### B. Decoding categorical neural responses using band frequency features and SVM

We investigated the decoding of prototypical from ambiguous vowels (i.e., category-level representations) using SVM neural classier on whole-brain (all 68 ROIs) and individual hemisphere (LH and RH) data separately for induced vs. evoked activity (FIG. 3 and Table I).

**FIG. 3.**
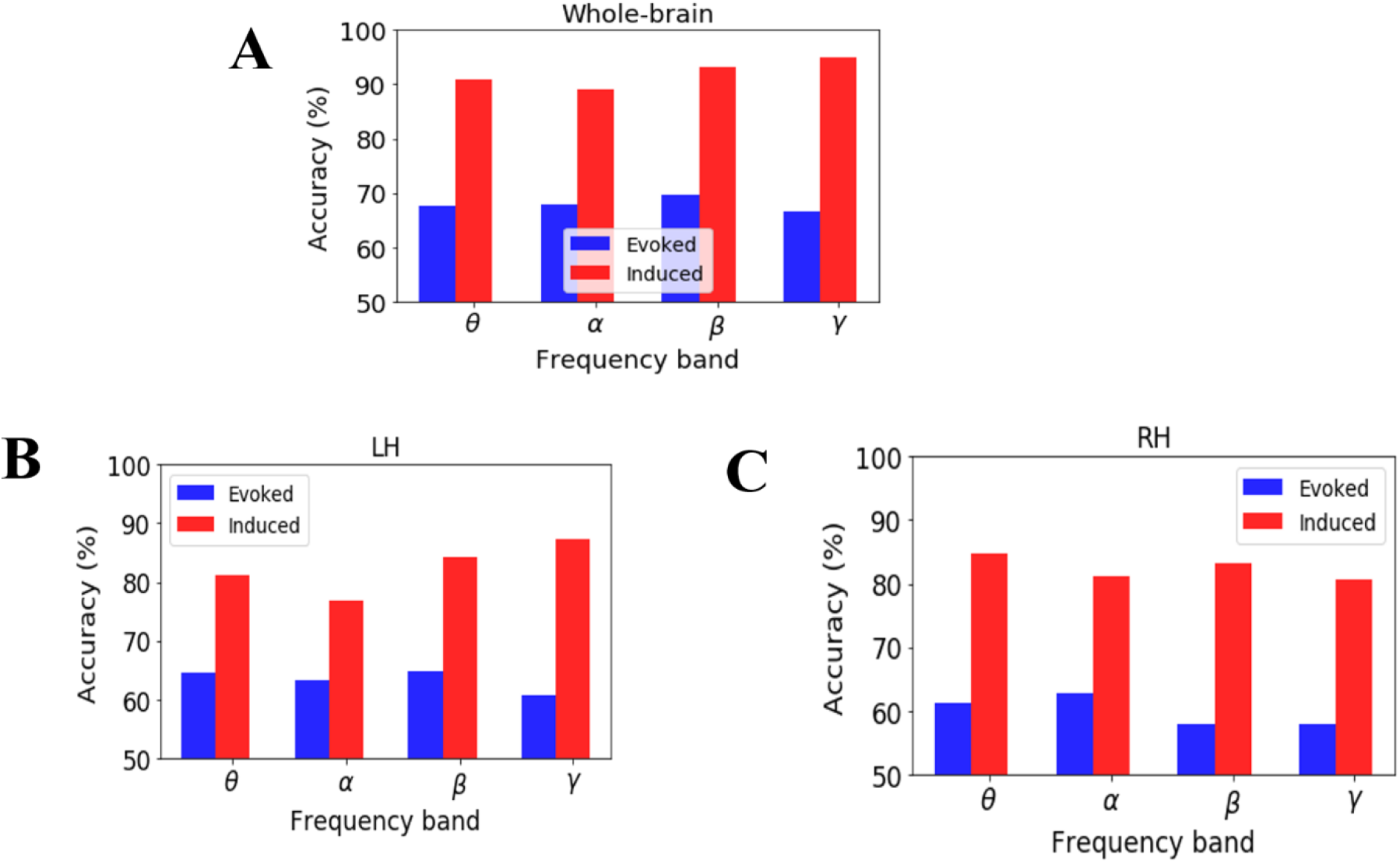
(Color online) Decoding categorical neural encoding using different frequency band features of source-level EEG. SVM results classifying prototypical (Tk1/5) vs. ambiguous (Tk 3) speech sounds. A) Whole-brain data (e.g. 68 ROIs), B) LH (e.g., 34 ROIs) C) RH (e.g., 34 ROIs). Change level =50%.

**Table I:** Performance metrics of the SVM classifier for decoding prototypical vs. ambiguous vowels (i.e., categorical neural responses) using whole-brain data.

| Neural activity | Frequency band | Accuracy<br>(%) | AUC<br>(%) | Precision<br>(%) | Recall<br>(%) | F1-score<br>(%) |
| --- | --- | --- | --- | --- | --- | --- |
| Evoked | $\theta$ | 67.53 | 67.59 | 68.00 | 68.00 | 67.00 |
| | $\alpha$ | 67.88 | 67.92 | 68.00 | 68.00 | 67.00 |
| | $\beta$ | 69.61 | 69.64 | 68.00 | 68.00 | 68.00 |
| | $\gamma$ | 66.66 | 66.65 | 67.00 | 67.00 | 67.00 |
| Induced | $\theta$ | 90.97 | 90.98 | 91.00 | 91.00 | 91.00 |
| | $\alpha$ | 89.06 | 89.05 | 89.00 | 89.00 | 89.00 |
| | $\beta$ | 93.23 | 93.20 | 93.00 | 93.00 | 93.00 |
| | $\gamma$ | 94.96 | 94.96 | 95.00 | 95.00 | 95.00 |

Using whole-brain *evoked* θ, α, β, and ɣ frequency responses, speech stimuli (e.g., Tk1/5 vs. Tk 3) were correctly distinguished at 66-69% accuracy. Among all evoked frequency bands, β-band was optimal to decode speech categories (69.61% accuracy). LH data revealed that θ, α, β, and ɣ bands decoded speech stimuli at accuracies between ~63-65% whereas decoding from RH was slightly poorer 57-62%.

Using whole-brain *induced* θ, α, β, and ɣ frequency responses speech stimuli were decodable at accuracies 89-95%. Among all induced frequency bands, ɣ band showed the best speech segregation (94.9% accuracy). Hemisphere specific data again showed lower accuracy. LH oscillations decoded speech categories at 76-87% accuracy whereas RH yielded 80-84%.

### C. Decoding brain regions associated with CP (evoked vs. induced)

We applied the stability selection (Meinshausen and Bühlmann, 2010) to induced and evoked activity features to identify the most critical brain areas (e.g., ROIs) that have been linked with speech categorization. Spectral features of brain ROIs were considered stable if the speech decoding accuracy was >70%. The effects of stability scores on speech sound classification is represented in FIG. 4. Each bin of the histogram illustrates the number of features in a range of stability scores. In this work, the number of features (labeled in FIG. 4) represents the neural activity of different frequency bands and the unique brain regions (labeled as ROIs in FIG. 4) represent the distinct functional brain regions of the DK atlas. The semi bell-shaped solid black and dotted red lines demonstrate classifier accuracy and area under the curve (AUC), respectively. We submitted the neural features identified at different stability thresholds to SVMs. This allowed us to determine whether the collection of neural measures identified via machine learning were relevant to classifying speech sound categorization.

**FIG. 4.**
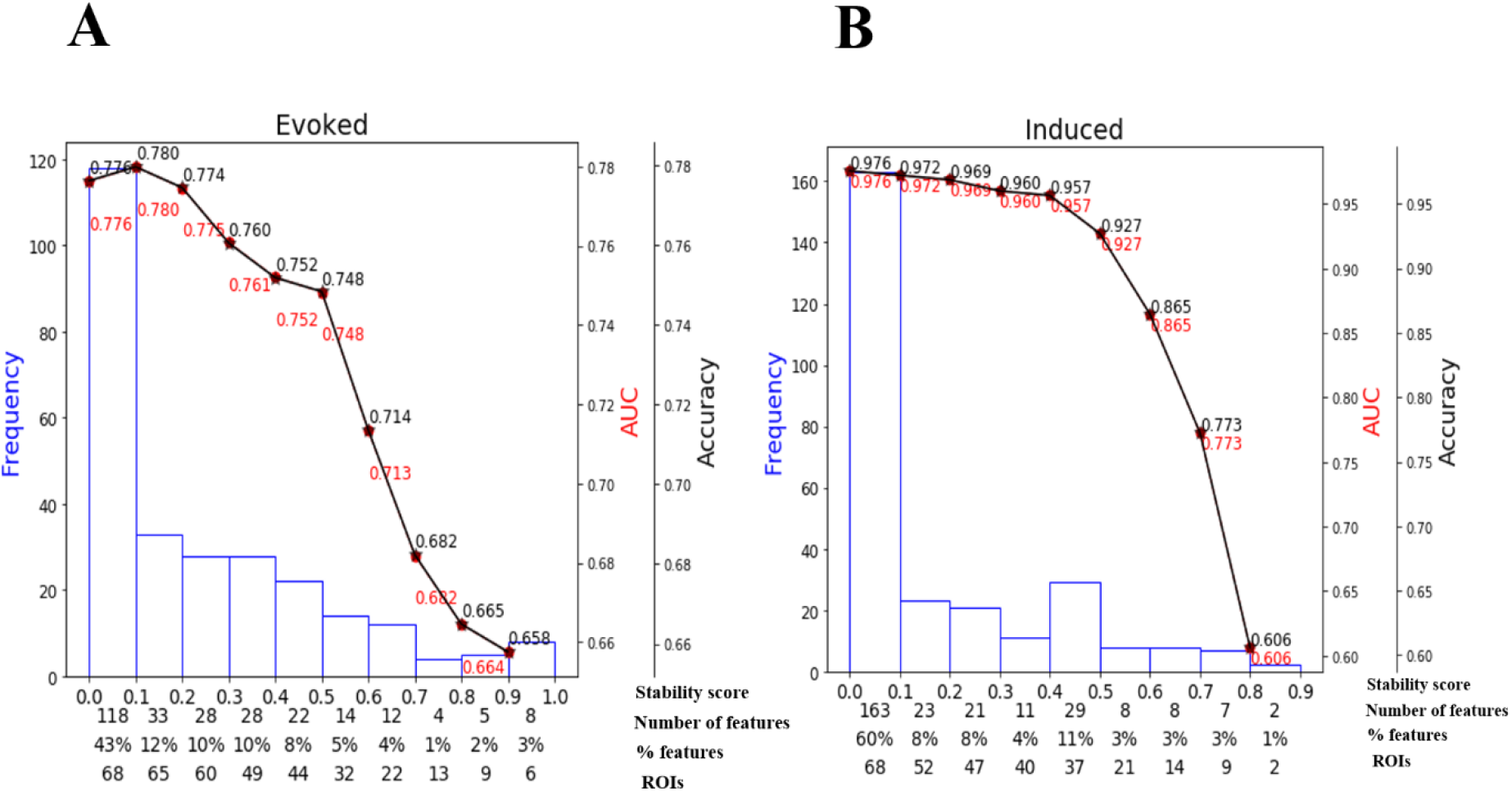
(Color online) Effect of stability score threshold on model performance during (A) evoked activity and (B) induced activity during CP task. The bottom of the x-axis has four labels; *Stability score* represents the stability score range of each bin (scores range: 0~1); *Number of features,* number of selected features under each bin; *% features,* the corresponding percentage of selected features; *ROIs*, number of cumulative unique brain regions up to the lower boundary of the bin.

For induced responses, most features (60%) yielded stability scores 0 to 0.1, meaning 163/272 (60%) were selected less than 10% out of 1000 iterations from 68 ROIs. A stability score of 0.2 selected 47/86 (32%) of the features from 47 ROIs that could decode speech categories at 96.9% accuracy. Decoding performance decreased with increasing the stability score (i.e., more conservative variable/brain ROIs selection) resulting in a reduced feature set that retained only the most meaningful features distinguishing speech categories from a few ROIs. For instance, corresponding to the stability threshold 0.5, 25 (10%) features were selected from 21 brain ROIs that yielded the speech categorization 92.7 % accurately. However, corresponding to the stability threshold 0.8 only 2 features were selected from 2 brain ROIs that decoded CP at 60.6%, still greater than chance level (i.e., 50%). Performance improved by ~10% (86.5%) when the stability score was changed from 0.7 (selected brain ROIs 9) to 0.6 (selected brain ROIs 14).

Using evoked activity, maximum decoding accuracy was 78.0% at a 0.1 stability threshold. Here, 43 % of features produced a stability score between 0.0 to 0.1. These 118 (43%) features are not informative because they decreased the model’s accuracy to properly categorize speech. Corresponding to the stability scores 0.9, only 8 features were selected from the 6 brain ROIs, which decoded speech at 65.8% accuracy. At stability score 0.6, 29 (1%) features were selected from 22 brain ROIs corresponding to 71.4 % accuracy performance. Thus, 0.6 might be considered an optimal stability score (i.e., knee point of a function in FIG. 4) as it decoded speech well above change (>70%) with a minimal (and therefore more interpretable) feature set for both induced and evoked activity. Classifier performance and brain ROIs corresponding to the optimal stability score (0.6) are shown in Table III in the Appendix.

### D. Brain-behavior relationships

To examine the behavioral relevance of the brain ROIs identified in stability selection, we conducted the multivariate weighted least square analysis (WLS) regression (Ruppert and Wand, 1994). We conducted WLS between the individual frequency band features (i.e., evoked and induced) and the slopes of the behavioral identification functions (i.e., FIG. 1B,) which indexes the strength of listeners’ CP. WLS regression for *induced* activity is shown in Table II (for evoked activity, see in the Appendix Table IV). From the induced data, we found that γ frequency activity from 6 ROIs predicted behavior best among all other frequency R^2^ = 0.915, p< 0.0001. Remarkably, only two brain regions (including PAC and Rostral anterior cingulate L) of β-band frequency could predict behavioral slopes (R^2^ =0.876, p<0.00001). Except in the α frequency band, evoked activity was poorer at predicting behavioral CP.

**Table II:** Brain-behavior relations of 14 brain ROIs in different frequency bands and behavioral prediction from the induced activity at a stability threshold ≥ 0.6 that yielded accuracy 86.5%.

| Frequency Band<br>and combined $R^2$ | ROI<br>ROI name | ROI<br>abbrev. | Coefficient | p-value |
| --- | --- | --- | --- | --- |
| $\theta$, $R^2 = 0.807$,<br>$p < 0.00001$ | Pars opercularis L | IPOP | 0.974 | 0.013 |
|  | Posterior cingulate L | IPCG | -1.759 | 0.001 |
|  | Caudal anterior cingulate R | rCAC | 3.163 | 0.001 |
| $\alpha$, $R^2 = 0.746$,<br>$p < 0.00001$ | Bankssts L | IBKS | 0.645 | 0.249 |
|  | Inferior parietal L | IIP | 1.594 | 0.035 |
|  | Precentral L | IPRC | 1.006 | 0.117 |
| $\beta$, $R^2 = 0.876$,<br>$p < 0.00001$ | Transverse temporal R | rTRANS | 0.267 | 0.584 |
|  | Rostral anterior cingulate L | IRAC | 3.004 | 0.001 |
| $\gamma$, $R^2 = 0.915$,<br>$p < 0.00001$ | Lateral occipital R | rLO | 0.768 | 0.804 |
|  | Lateral orbitofrontal R | rLOF | 5.092 | 0.001 |
|  | Superior parietal R | rSP | 16.472 | 0.004 |
|  | Caudal middle frontal R | rCMF | -3.243 | 0.188 |
|  | cuneus R | rCUN | -1.743 | 0.701 |
|  | Lateral orbitofrontal L | ILOF | 0.709 | 0.553 |

**Table III:**
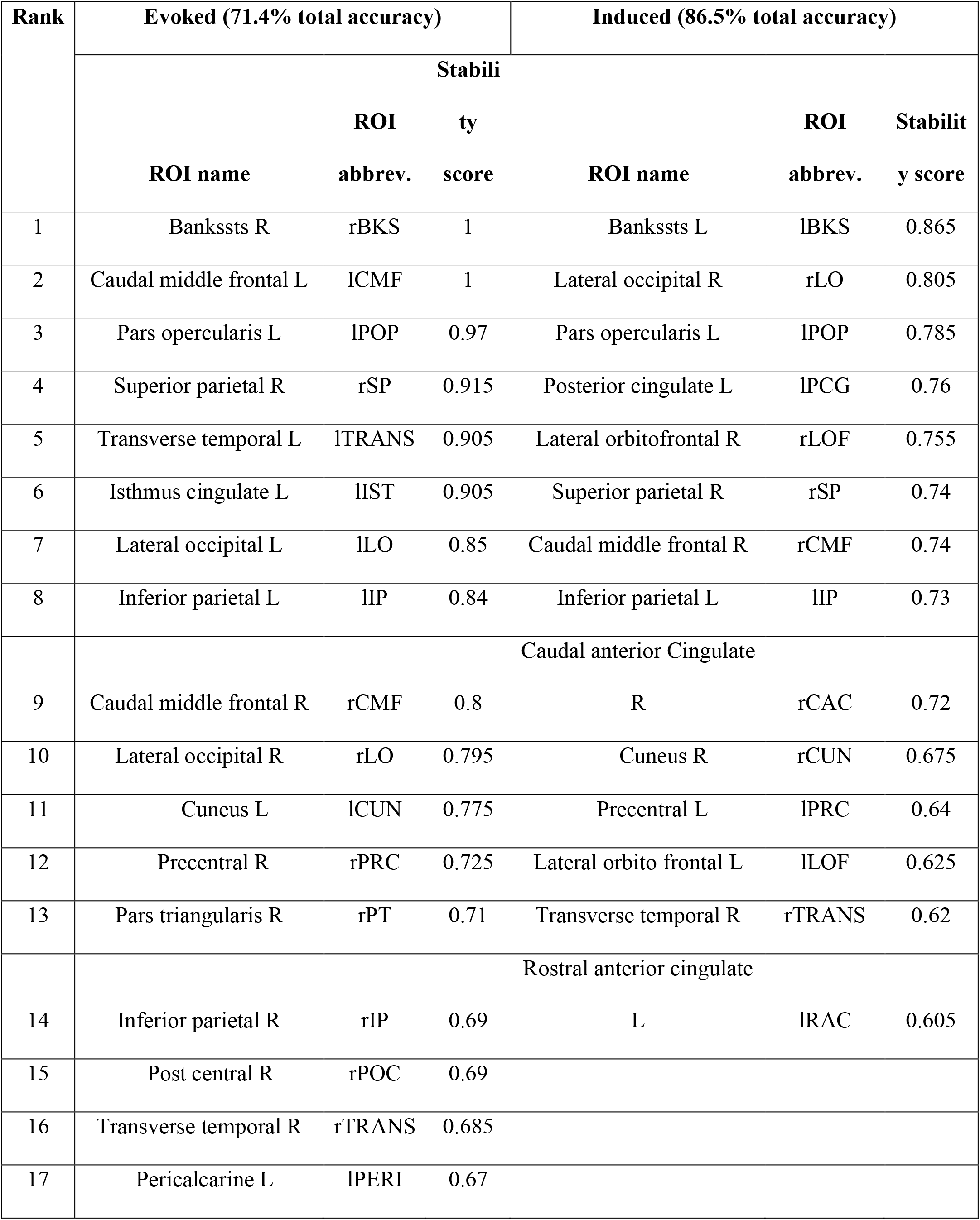
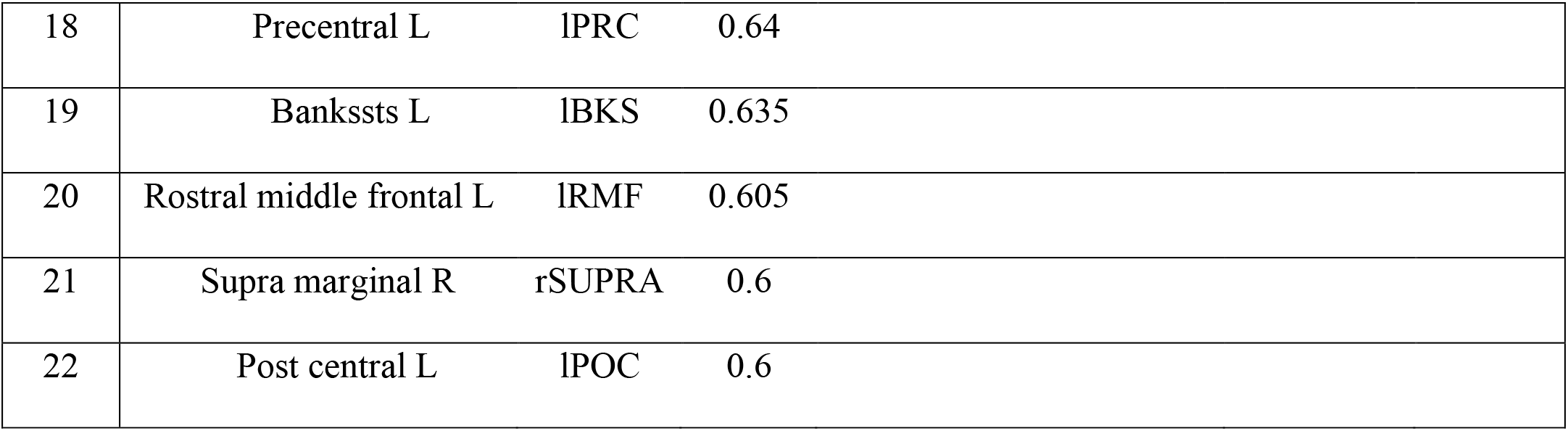
Most important brain regions describing speech categorization. The *evoked activity* identified (22 ROIs) and *induced activity* identified 14 ROIs at a stability threshold ≥ 0.6. Corresponding to this threshold evoked and induced activity showed 71.4%, 86.5% accuracy respectively.

**Table IV:**
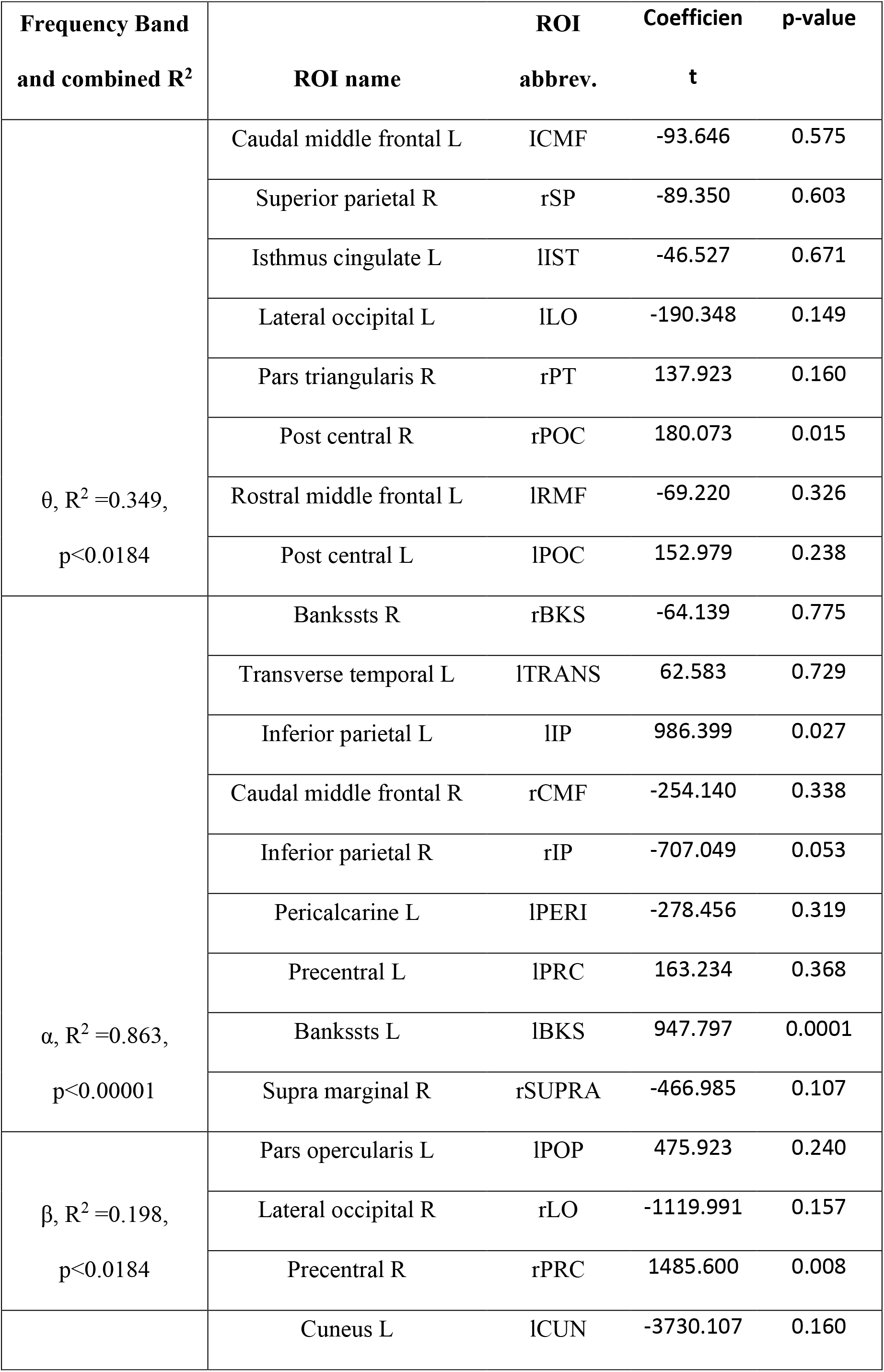
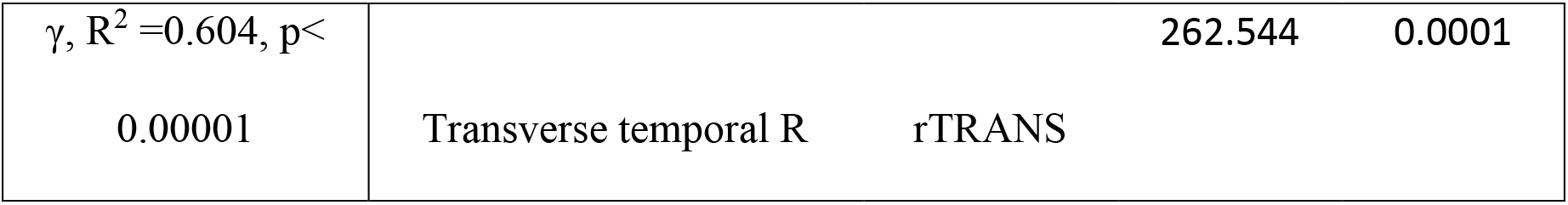
Brain-behavior relations of 22 brain ROIs in different frequency bands and behavioral slope prediction from the evoked activity at a stability threshold ≥ 0.6 that yielded accuracy 71.4%.

## IV. DISCUSSION

### A. Speech categorization from evoked and induced activity

The present study aimed to examine which modes of brain activity and frequency bands of the EEG best decode speech categories and the process of categorization. Our results demonstrate that at the whole-brain level, evoked β-band oscillations robustly code (~70% accuracy) category structure of speech sounds. However, induced ɣ-band showed better performance, classifying speech categories at ~95% accuracy, better than all other induced frequency bands. Our data are consistent with notions that higher frequency bands are associated with speech identification accuracy and carry information related to acoustic features and quality of speech representation (Yellamsetty and Bidelman, 2018). Our results also corroborate previous studies that suggest higher frequency channels of the EEG (β, γ) reflect auditory perceptual object construction (Tallon-Baudry and Bertrand, 1999) and how well listeners map sounds to category labels (Bidelman, 2015, 2017).

Analysis by hemispheres showed that induced γ activity was dominant in LH whereas lower frequency band (e.g., θ) were more dominant in RH. These findings support the asymmetric engagement of frequency bands during syllable processing (Giraud et al., 2007; Morillon et al., 2012) and lower frequency band in RH dominance in inhibitory and attentional control (top-down processing during complex tasks) (Garavan et al., 1999; Price et al., 2019). Our results are consistent with the idea that cortical theta and gamma frequency bands play a key role in speech encoding (Hyafil et al., 2015). They also show that the machine learning model was able to decode acoustic-phonetic information (i.e., speech categories) in LH (using induced high frequency) and task difficulty (using low frequencies) in RH.

### B. Brain networks involved in speech categorization

Machine learning (stability selection coupled with SVM) further identified the most stable, relevant, and invariant brain regions that associate with speech categorization. Our results show that induced activity better exhibited speech categorization using less neural resources (i.e., minimum brain ROIs) as compared to evoked activity. Using induced activity, stability selection identified 14 critical brain regions including the primary auditory cortex (Transverse temporal R), Brocas’s area (Pars opercularis L), and motor area (Precentral L). These ROIs are widely engaged in speech-language processing. Superior parietal and inferior parietal areas have been associated with auditory, phoneme, and sound categorization in particularly ambiguous contexts (Dufor et al., 2007; Feng et al., 2018). The orbitofrontal is associated with speech comprehension and rostral anterior cingulate with speech control (Sabri et al., 2008). Surprisingly, out of the identified 14 brain ROIs; three ROIs are in θ, three in α, two in β, and six in γ band. Noticeably, we found that a greater number of brain regions were recruited in the γ-frequency band. This result is consistent with the notion that high-frequency oscillations play a role in network synchronization and widespread construction of perceptual objects related to abstract speech categories (Giraud and Poeppel, 2012; Haenschel et al., 2000; Si et al., 2017; Tallon-Baudry and Bertrand, 1999). Indeed, γ-band activity in only six ROIs were the best predictor of listeners’ behavioral speech categorization.

In sum, our data suggest that induced neural activity plays a more prominent role in describing the perceptual-cognitive process of speech categorization than evoked modes of brain activity (Doelling et al., 2019). In particular, we demonstrate that among these two prominent functional modes and frequency channels characterizing the EEG, induced γ-frequency oscillations best decode the category structure of speech and the strength of listeners’ behavioral identification. In contrast, the evoked activity provides a reliable though weaker correspondence with behavior in all but the α frequency band.

## ACKNOWLEDGMENTS

This work was supported by the National Institutes of Health (NIH/NIDCD R01DC016267) and the Department of Electrical and Computer Engineering at the University of Memphis. Requests for data and materials should be directed to G.M.B.

## APPENDIX

## SVM optimal parameters values

The optimal parameters of SVM classifier are given below in different analysis scenarios. For induced activity, the optimal values were used for full brain data: [C=10, ɣ=0.001 for θ-band, C=20, ɣ=0.001 for α-band, C=40, ɣ=0.001 for β-band, C=30, ɣ=0.001 for γ-band]; LH data: [C=10, ɣ=0.001 for θ-band, C=20, ɣ=0.001 for α-band, C=40, ɣ=0.001 for β-band, C=30, ɣ=0.001 for γ-band]; RH data:[C=10, ɣ=0.001 for θ-band, C=20, ɣ=0.001 for α-band, C=40, ɣ=0.001 for β-band, C=30, ɣ=0.001 for γ-band].

For evoked activity, the optimal values for the full brain data are: [C=30, ɣ=0.001 for θ-band, C=40, ɣ=0.0001 for α-band, C=40, ɣ=0.001 for β-band, C=10, ɣ=0.0001 for γ-band]; LH data: [C=20, ɣ=0.001 for θ-band, C=20, ɣ=0.001 for α-band, C=30, ɣ=0.002 for β-band, C=30, ɣ=0.003 for γ-band]; RH data:[C=10, ɣ=0.004 for θ-band, C=20, ɣ=0.003 for α-band, C=20, ɣ=0.002 for β-band, C=20, ɣ=0.002 for γ-band].

## Stability selection

Stability selection is a state-of-the-art feature selection method that works well in high dimensional or sparse data based on the Lasso (least absolute shrinkage and selection operator) (Meinshausen and Bühlmann, 2010; Yin et al., 2017). Stability selection can identify the most stable (relevant) features out of a large number of features over a range of model parameters, even if the necessary conditions required for the original Lasso method are violated (Meinshausen and Bühlmann, 2010).

In stability selection, a feature is considered to be more stable if it is more frequently selected over repeated subsampling of the data (Nogueira et al., 2017). Basically, the Randomized Lasso randomly subsamples the training data and fits a L1 penalized logistic regression model to optimize the error. Over many iterations, feature scores are (re)calculated. The features are shrunk to zero by multiplying the features’ co-efficient by zero while the stability score is lower. Surviving non-zero features are considered important variables for classification. Detailed interpretation and mathematical equations of stability selection are explained in (Meinshausen and Bühlmann, 2010). The stability selection solution is less affected by the choice of the initial regularization parameters. Consequently, it is extremely general and widely used in high dimensional data even when the noise level is unknown.

